# A Continuous, Low-Flow, and Multiplexing Pumping System for Microfluidics Applications

**DOI:** 10.1101/2024.08.16.608339

**Authors:** Kara A. Walp, Yash H. Patel, Woud Alsadoun, Hailey Lenn Gordon, Hamed Rastaghi, Samuel M. D. Oliveira

## Abstract

Microfluidics optimize experimental procedures but often require external pumps for precise, steady, and low flow rates. These procedures typically require extended, continuous operation for long-duration experiments. We introduce the Dual-Syringe Continuous Pumping Mechanism (DSCPM), a low-cost, precise, and continuous pump for microfluidic applications with input multiplexing capability. With a 3D-printed housing and standard components, the DSCPM is easy to fabricate and accessible. Operating at a microliter per minute flow rates, the DSCPM uses fluidic bridge rectification to combine syringe pump precision with continuous infusion. We validated laminar flow in microfluidic ’cell traps’ without disrupting microbial growth. COMSOL simulations confirmed safe shear stress levels. We also developed and tested fluidic multiplexers for greater modularity and automation. Addressing current pump limitations, such as discontinuity and high costs, the DSCPM can enhance experimental capabilities and promote efficiency and precision while increasing accessibility of hardware automation in many fields.

## Introduction

Microfluidics is critical in many biological research fields because it allows for automation and high-throughput experimentation. It is leveraged to reduce system footprint, conduct low-volume experiments, and more. Many systems operate for extended experimental durations and require external infusion devices for low flow rates. Current research, including endothelial microfluidics and biofilm studies, underscores these requirements, calling for long-term flow generation in the microliter per minute regime, as low as 1.67 μL/min^1–3^. Microfluidic chips with a cell trapping geometry called monolayer chambers (MCs), which are used for microbial housing and characterization; for example, for prototyping a synthetic gene circuit^4^; allow for continuous facilitation of log phase proliferation of microbes by washing away excess cells and continuously supplying media^5–7^. This introduces the need for constant flow generation at low flow rates^6^ that is steady enough to avoid harmful shear stress on housed cells. In addition to continuity and precision, the need for automation of flow input perturbations is demonstrated in synthetic biology for synthetic microbial consortia with cascade-inducing inputs, among other applications^4,8–14^.

Various existing pumping mechanisms, such as peristaltic pumps, pressure/diaphragm pumps, and syringe pumps (Figure 1) can continuously infuse into a system from a reservoir. Still, they typically lack the resolution to generate steady flow at low flow rates. Peristaltic pumps enclose flexible tubing in a rigid housing and compress the tubing cyclically to generate flow. The regime of flow rates reachable with peristalsis depends on the resolution of the compression system and the inner diameter (ID) of the tubing. Existing peristaltic micropumps report achievable flow rates as low as 50μL/min (Taskgo Fluidics). Alternative solutions using PDMS inlays report small volume liquid handling but does not report minimum flow rates^15^. Another peristaltic pump study reports flow rates sub 1 μL/min. However, the metrics determining the minimum flow rate used in that study were undisclosed.^16^ Additionally, the presented design reported a sudden change in flow rate at the point of release of roller pins^16^, which is a critical flaw in the applications this study aims to accommodate-large fluctuations in flow rate from a pumping mechanism introduce potentially harmful shear stress. Additionally, peristaltic pump tubing can be replaced to shift the regime of reachable flow rates for a given pump. However, the diameter of flexible tubing can only get so small before the material is no longer structurally sound. These parameters constrain the minimum flow rates of conventional peristaltic pumps to higher than those required for some micro/milli-fluidic applications, including infusion into microfluidics which contain cell trapping MCs. On the other hand, pressure pumping systems modulate flow generation using pressure sensing and feedback mechanisms, meaning that the system is prone to oscillations, and precision is difficult to obtain. Most small pressure/diaphragm pumps operate in a larger order of magnitude than is intended in this study, like a piezoelectric diaphragm pump which was designed by Taskago Fluidics which can reach flow rates as low as 3mL/min. Another piezoelectric micropump, closer to our intended range, was cited to operate <3.14 μL/min, but it was integrated into a microfluidic device and required expensive equipment to fabricate^17,18^. An electrostatic micropump was designed to generate flow in the 30 μL/min regime, but requires inaccessible operative voltages (160V)^19^. Additionally, previous studies report pressure-actuated acrylic valves able to pump in the regime of <20μL/min but were not able to provide any flow rate variance or laminarity characteristics and required access to CNC manufacturing machinery, which limits the accessibility of pump fabrication^20^.

**Figure 1.**
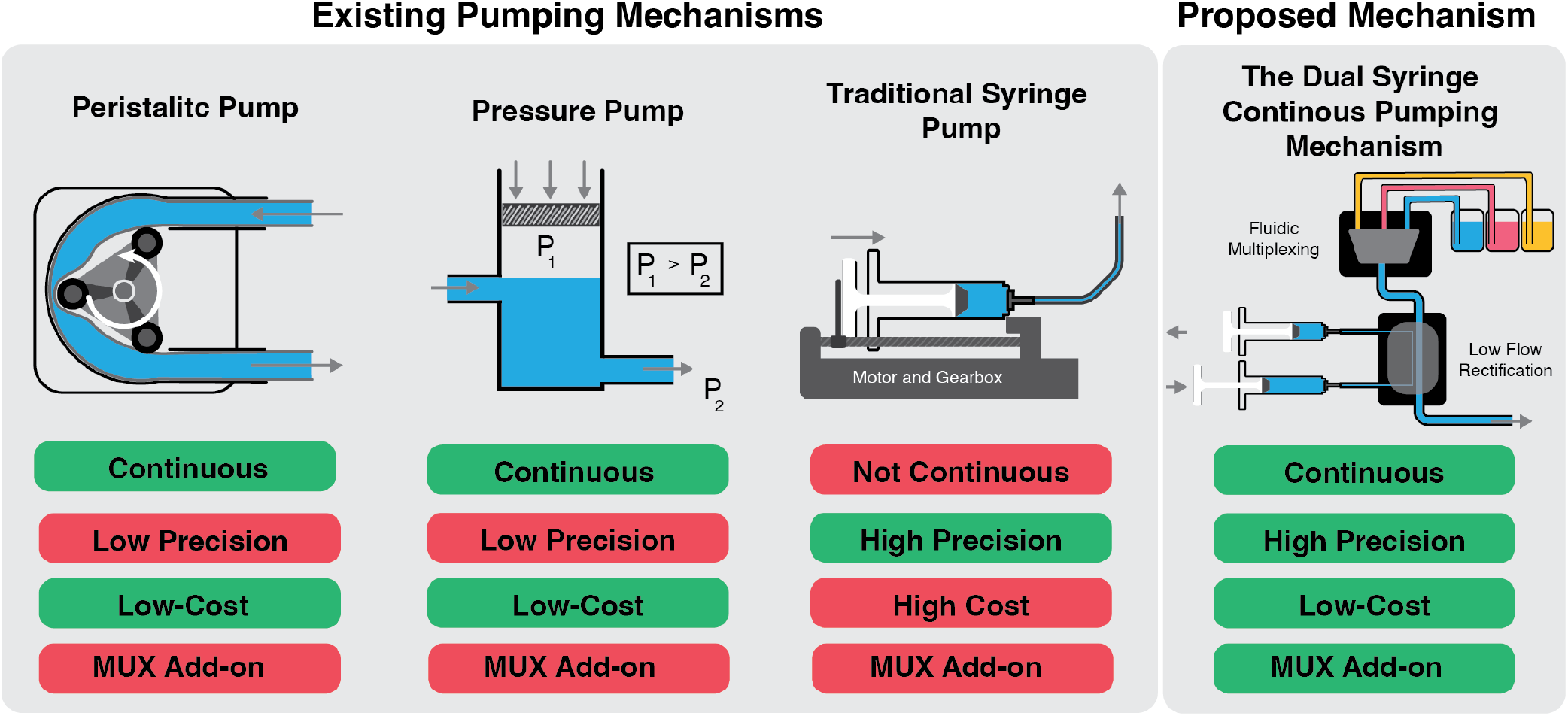
Comparison of established pumping technologies (peristaltic pump, pressure pump, and traditional syringe pump) to the proposed technology, the dual syringe continuous pumping mechanism (DSCPM), in the scope of biological and microfluidic applications. This work presents a precise, low-cost solution for complex microfluidic pumping applications.

Conventionally, syringe pumps are used for low flow rate applications (<100μL/min). Syringe pumps use motor-driven pushers to manipulate the plunger of a syringe. The minimum sufficiently laminar flow rate they can reach is determined by two parameters-the linear actuation system and the inner diameter of the syringe: this level of direct control of flow generation results in a highly precise pumping mechanism. A syringe (or “mechanical”) micropump that used piezoelectric linear actuation was cited to produce stable flow at 4 nL/min, with the volume infused by a single pump stroke being 511 nL^21^. However, traditional bench-top syringe pumps are often expensive, large, and are considered a gold-standard in low-flow rate microfluidics. While seemingly ideal for micro-scale experimentation, one significant drawback to using traditional syringe pumps is that they are not continuous since they require the syringe to be manually refilled once its total initial volume has been infused. This can be detrimental when experiments must be run for longer durations, since manually refilling the syringe every time it depletes is not always feasible.

When designing our pumping system, we combined the high-precision flow generation mechanism of syringe pumping with the electronics principle of bridge-rectification to achieve continuous flow. Our system uses two syringes that infuse and refill inversely to one another, while three-way solenoid pinch valves manually rectify the flow. In other words, as one syringe infuses, the second syringe refills; they switch directions periodically, and valves direct the generated flow forward. Rectifying dual-syringe produced flow to build a continuous infusion system has been done before (See New Era Pump Systems DUAL-NE-1000X Continuous Infusion Syringe Pump System However, each market-available solution that employs this concept uses check valves for flow rectification, which disrupt flow laminarity at low flow rates (< about 100μL/min). Another pitfall is that each existing solution consists of a “kit” of rectifying equipment meant to be installed on traditional syringe pumps, meaning that one continuous infusion unit costs thousands of dollars due to the market value of conventional syringe pumps.

Here, we present an open-source, low-cost, dual-syringe continuous pumping mechanism (DSCPM) with seamlessly integrable fluidic multiplexers. Our system combines the high-precision flow generation mechanism of syringe pumping with the electronics principle of bridge-rectification to achieve continuous flow, using two syringes that infuse and refill inversely to one another, while three-way solenoid pinch valves manually rectify the flow. The DSCPM consists of a 3D printable housing (Figure S6), various low-cost electronics components, such as a servo motor and an Arduino Uno Micro-controller (Figure S1), and various low-cost fluidics components (such as syringes, Luer locks, and silicone tubing). The DSCPM costs approximately $500 for all components (Table S1). The DSCPM can reach flow generation precision that rivals existing traditional syringe pumps. Due to the nature of the flow rectification mechanism that the DSCPM employs, the DSCPM infuses directly from an external reservoir, meaning that it is completely continuous and could infuse theoretically indefinitely, given that the reservoir has a large enough volume. The easily integrable fluidic multiplexers allow for perturbations between and mixing multiple sources and consist of three way solenoid pinch valves, a micro-controller, and additional tubing.

## Results

### The Dual Syringe Continuous Pumping Mechanism Design

#### Flow Generation

With traditional syringe pumps, volume is a constraint since once a syringe is depleted, it must be manually refilled. With dual syringe systems that have automated flow rectification, this is no longer a constraint, meaning much smaller syringes can be used, and less complexity of the linear actuation mechanism is required to generate the same volumetric flow rates.

Syringes of less volume tend to have smaller inner diameters. Syringe cross sectional area can be defined by:

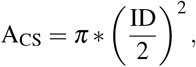

where A_CS_ is the cross-sectional area, and ID is the Syringe’s Inner Diameter

The cross-sectional area is directly proportional to the volume infused per minimum motor increment, which can be defined by:

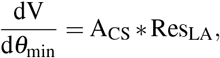

where 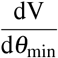 is the minimum volume increment achievable, Res_LA_ is the linear resolution of the Linear Actuator

The DSCPM prototype uses 10 μL syringes. Theoretically, the prototype can produce laminar flow at flow rates as small as 0.152 μL/min. The calculation of the minimum laminar flow rate from the volume infused per minimum motor increment was derived from the specifications of the Harvard Apparatus PhD 2000 Syringe Pump. In the PhD 2000’s case, the motor takes 12,800 μ-steps per revolution with 1/16th micro-stepping, with a distance between grooves on the lead screw yielding 1.05 mm of linear motion per revolution. This gives us:

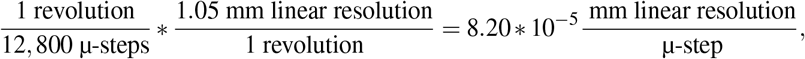

By reverse engineering published specifications for the Harvard Apparatus PhD 2000 Syringe Pump, we were able to derive a metric that defines the minimum flow rate a pump can generate with adequate stability, or “laminarity”, which was a maximal duration between minimum motor increments of 27.34 seconds (Supplementary Methods 1).

Using this metric, the DSCPM prototype (which yields 10 μL syringes) can theoretically generate stable flow at 14.0 nL/min. The smallest syringes that the DSCPM could employ are 0.5 μL syringes; this setup could generate stable flow at 7.0 nL/min (Figure S7). Based on these calculations and metrics additional flow regimes can also be achieved (Table S2).

#### Flow Rectification

The DSCPM generates flow using two syringes that simultaneously infuse and refill. This can be equated to alternating current (AC) voltage. AC voltage must be rectified in electronics to produce direct current (DC) voltage, usually using bridge rectifiers. This principle can also be applied in fluidics^22,23^. The means of rectification typically employed by existing dual syringe systems is check valves, which are the fluidic equivalent of diodes. Check valves usually have an internal spring-loaded mechanism, only allowing flow in one direction (Figure 2B). If check valves behaved ideally, they would pose a perfect rectification tool; however, similarly to how electronic diodes have a threshold forward voltage that must be reached before current can flow, check valves have a minimum cracking pressure that must be reached before the valve “opens” and allows forward flow (See Figure 2C). At most flow rates, the effect of the minimum cracking pressure on the rectified flow is negligible; however, at flow rates in the regime of 100s of μL/min and below, laminarity was observed to be detrimentally damaged by check valves.

**Figure 2.**
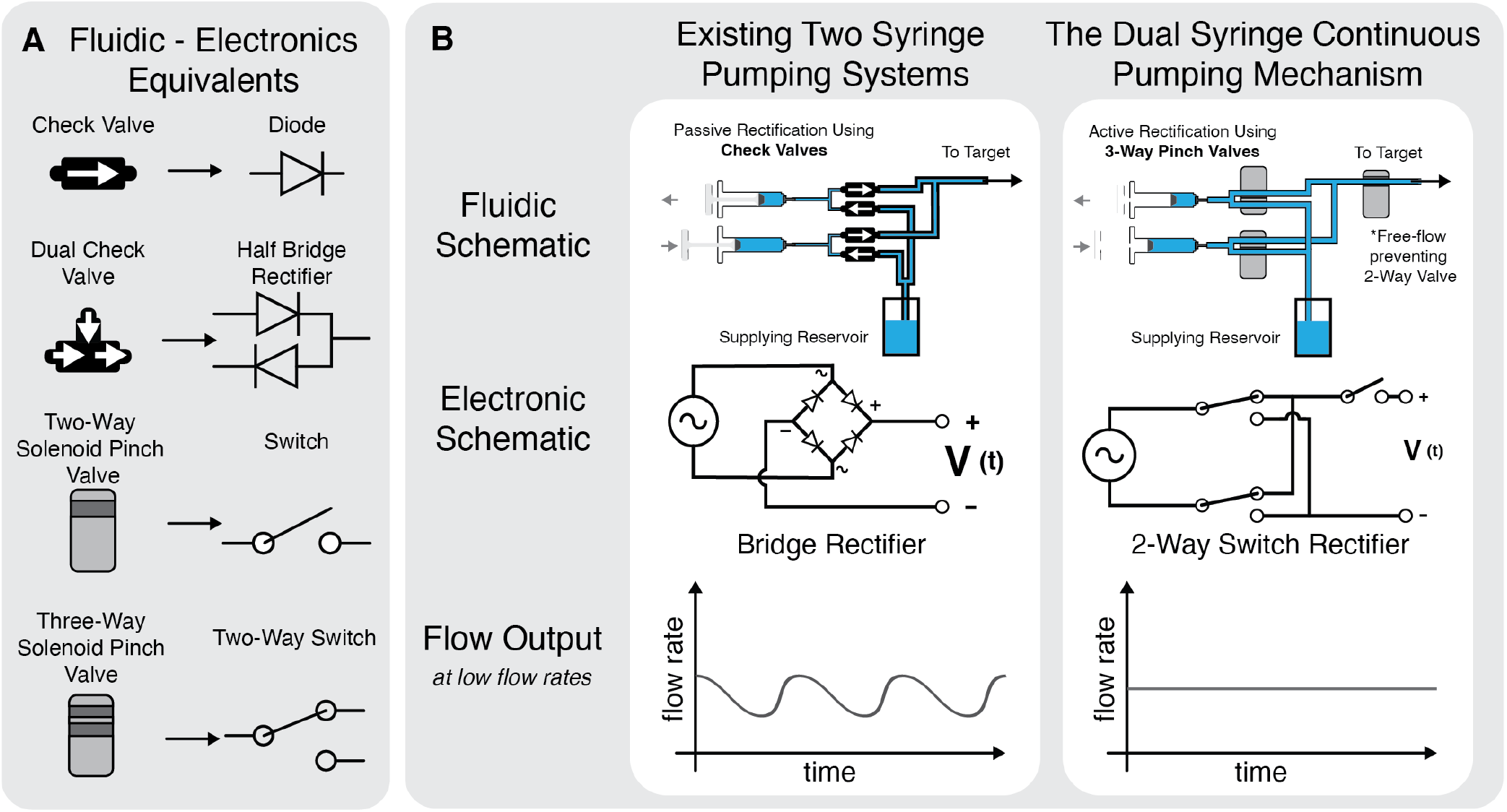
Development of the Dual Syringe Continuous Pumping Mechanism from Electronics Principles. A) A library of fluidic/electronic equivalents. B) Comparison of existing two syringe pumping systems for passively rectifying flow with check valves and the dual syringe continuous pumping mechanism, which rectifies with solenoid pinch valves. Along with their expected flow output, which is informed by the behavior of their electrical equivalent.

It is hypothesized that when using valves with considerable cracking pressures for low flow applications, due to external factors including the elasticity of most fluidics components, and intrinsic inductive properties of most check valves^24^ (inertia delaying valve close time), the time it takes to build up to the minimum cracking pressure is large enough to render the flow no longer laminar, and essentially results in periodic bolus infusions instead of laminar flow (See Figure 2A). This means that even if the flow is generated with very sophisticated syringe pumps with highly precise linear actuation mechanisms, using check valves to rectify the flow will defeat the purpose of using such pumps. Additionally, Check valves with lower cracking pressures were observed to allow small amounts of backflow in the case of a negative pressure gradient across the valves. A study that designed and fabricated microfluidic check valves specifically for flow rectification cited a maximum rectification efficiency of 29.8%^25^. This is also true of other passive, non spring loaded valve types, such as microfluidic diodes, which do not exhibit 100% efficiency and are subpar for low Reynolds number applications^23^.

#### DSCPM Schematics

A logical fallacy was observed affecting flow behavior when using the rectification schematic that employs two three-way pinch valves (called the DSCPM 2-3 Schematic). With the DSCPM 2-3 schematic, the three-way valves switch states simultaneously once per syringe cycle. With a three-way valve, when the valve pusher is mid-switch, there is an instant where neither tubing branch is completely pinched, and free flow is allowed between the three branches of tubing controlled by the valve (See the “danger zones” in Figure 3B). The time it takes for a three-way valve to leave the “danger zone” when it switches states is on the scale of milliseconds; it was initially anticipated to be instantaneous enough to not affect the flow generated by the DSCPM. Unfortunately, at low flow rates (<50μL/min), the free flow during the “danger zones” was large enough to significantly affect the overall flow. This phenomenon is equivalent to if an electronic three-way switch instantaneously short-circuited its three terminals while switching states, instead of instantaneously opening the circuit.

**Figure 3.**
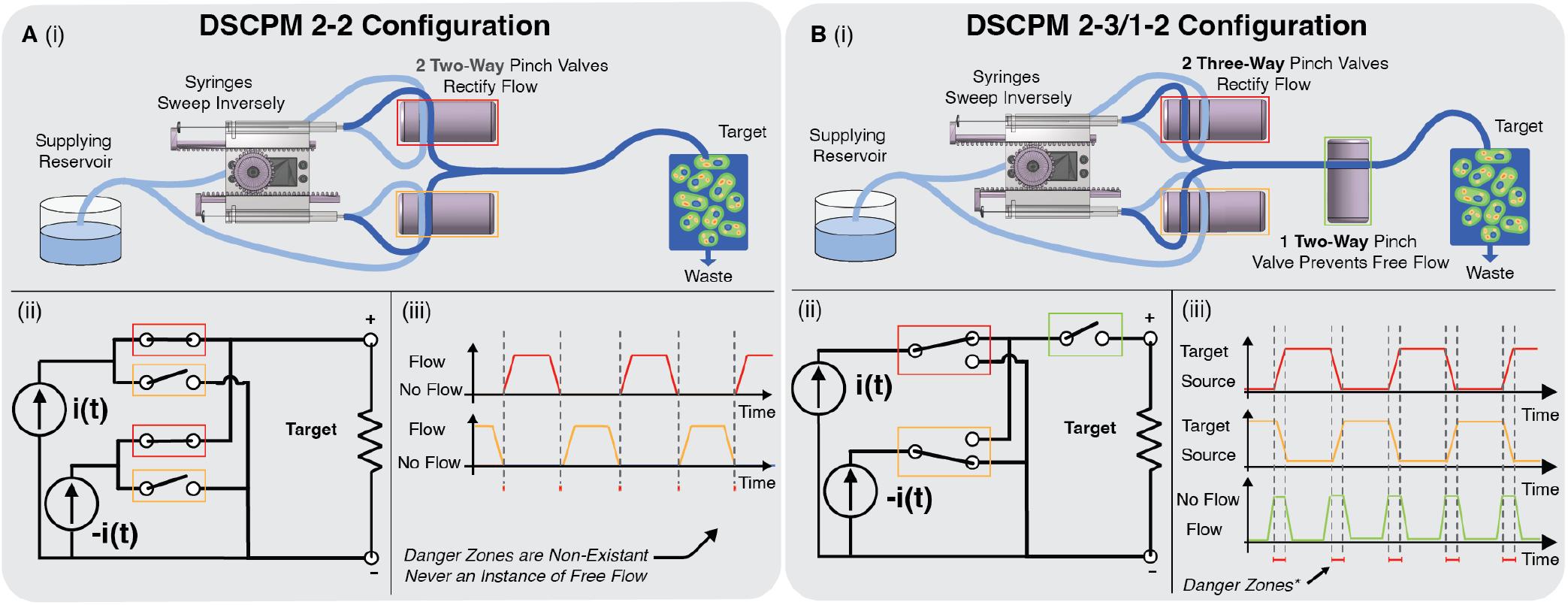
DSCPM “wiring diagram” Schematics. A) i) Schematic of the DSCPM 2-2, which signifies use of two 2-way pinch valves. ii) The electronic equivalent of the DSCPM 2-2. iii) Waveform describing the behavior of the pinch valves while the DSCPM 2-2 is infusing. B) i) Schematic of the DSCPM 2-3/1-2, which signifies use of two 3-way pinch valves and one 2-way pinch valve. ii) The electronic equivalent of the DSCPM 2-3/1-2. iii) Waveform describing the behavior of the pinch valves while the DSCPM 2-3/1-2 is infusing.

To eliminate the possibility of any free flow during the “danger zones,” two new DSCPM configurations were conceptualized. The DSCPM 2-3/1-2 (See Figure 3B) refers to a configuration with two three-way valves and one two-way valve, and the DSCPM 2-2 refers to a configuration with two two-way valves (See Figure 3A). The DSCPM 2/3-1/2 is the most straightforward solution and a logical continuation from the DSCPM 2-3 which had unprotected flow; a two-way, normally open pinch valve is added in series with the rectification logic, and pulses on (stops all flow) in the instant that the two rectifying valves switch states (the danger zone). The alternative schematic, the DSCPM 2-2, requires a slightly different tubing setup that has two cross-sections of tube per two-way valve and has slightly staggered state switching so that there is never an instant where both valves are open.

A consequence of both solutions is the introduction of a momentary flow rate spike that occurs periodically as the valves cycle. When pinch valves pinch tubing, the fluid in the pinched cross-section is displaced rapidly. This phenomenon did not cause issues with the original, unprotected schematic (the DSCPM 2-3) since all valves switched states simultaneously; equal amounts of cross-sections were pinched and un-pinched, rendering the sum of displaced fluid volume zero. With the DSCPM 2-3/1-2, the tubing cross-section fitted with the free-flow-preventing two-way valve is unaccounted for by complimentary tubing; this results in one flow rate spike per cycle induced by one tubing cross-section of volume. With the DSCPM 2-2, the staggered state switching means that the pulse will be induced by the volume of two tubing cross-sections, and will theoretically be twice the size of the DSCPM 2-3/1-2 pulse. While the DSCPM 2-2 is more monetarily efficient (two valves instead of three valves), the DSCPM 2-3/1-2 has pulses that are half the magnitude of the DSCPM 2-2 pulses and is, therefore, more optimal in terms of flow rate consistency and laminarity. Due to its superiority in terms of precision, the DSCPM 2-3/1-2 will be the schematic used for all laminarity characterization and is the recommended schematic to implement.

### Quantitative Characterization of Infusion at Low Flow Rates

To characterize the flow generated by the DSCPM, volume infused over time was measured and derived into flow rate over time for three small flow rates (1.5, 10, and 25 μL/min). The DSCPM was configured to infuse dyed water into micro-bore tubing (0.030” ID), and 150 mm of the tubing was visually tracked (filmed)^26^ as the DSCPM infused into it. The footage was cropped and processed with an algorithm that utilizes an edge detection kernel to quantify fluid position as a function of time. Using the tubing dimensions, volume infused as a function of time was calculated, from which the volumetric flow rate was derived.

This approach was also used to characterize the flow direction reversal response time of the DSCPM. As microfluidics advance, the need for complex flow behavior is emerging. In traditional syringe pumps with advanced linear actuation that use large syringes, flow direction reversal time (the time it takes for flow to reverse once it is commanded) was observed to be very large, given that the syringes generally used have plastic pushers which are slightly elastomeric, which requires syringe priming upon direction reversal (https://www.syringepump.com/continuous.php). The 10 μL syringes used in the DSCPM prototype are glass with steel pushers, which are not compressible, meaning that direction reversal is near instant and should not require syringe priming.

To further analyze the behavior of the DSCPM flow, computational fluid dynamics (CFD through COMSOL Multiphysics V6.1 with the Microfluidics Module) were used to simulate one pressure spike inside of a cell trapping MC, which is a type of cell trapping microfluidic geometry. A metric of laminarity is often defined by the Reynolds number (Re), a dimensionless measurement of the viscous forces and inertial forces present in a flow system defined by: Re 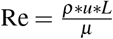 where, *μ* is dynamic viscosity, *ρ* is density, *u* is flow velocity, and *L* is characteristic length. Flow is considered laminar when Re is between 0 and 2320; anything greater than 4000 is considered Turbulent^27^. A simulation assuming laminar flow in 3D channels were carried out to determine laminarity properties given our measured system^28^. Based on flow results, the calculated Re for each flow rate is, on average, 9.488, 4.498, and 0.6787 for flow rates of 25, 10, and 1.5 μL/min. The impact of values results in a spike in flow rate, which results in a variation of the Re in the range of 15.053 - 24.495, 8.972 - 23.674, and 5.068 - 1.368 for 25, 10, and 1.5 μL/min respectively.

The DSCPM’s flow at low flow rates was also characterized and compared to the industry gold standard of high-precision flow by measuring relative pressure over time. A Honeywell ABP2 Series Differential Pressure Sensor, configured as a monometer, was linked in series with the DSCPM prototype (Figure S2). Water was infused into a microfluidic chip with cell trapping MCs at three different flow rates (1.5, 10, and 25 μL/min). These measurements were replicated using a Harvard Apparatus PhD 2000 syringe pump to compare values. The standard deviations of the DSCPM series’ relative pressures were 0.0395, 0.1192, and 0.0745 PSI for the respective flow rates. For the HA PhD 2000, the values were 0.0219, 0.0263, and 0.0290 PSI. The pressure magnitudes at the flow rate spikes were then isolated and quantified, yielding values of 0.7708 ± 0.3836, 0.6913 ± 0.2598, and 0.4411 ± 0.4351 PSI for the respective flow rates.

### Long-Term Maintenance of a Microbial Consortia

In research involving microbes, cell trapping MCs are commonly used to house microbes. In the case of bacterial cells, MCs are helpful because the flow of media past an MC of the right dimensions can ensure that the contained microbes get proper nutrition and that once the microbes have grown to fill the MC, any excess microbes are washed away in the flow of media. This is a prime use case for the DSCPM, given that high precision and continuous flow is needed.

The DSCPM was used to infuse media at 1.5 μL/min into a polydimethylsiloxane (PDMS) microfluidic chip with MCs containing microbes genetically engineered (see Methods) to constitutively express cyan fluorescent protein (CFP). The experiment lasted 24 hours, and growth and proliferation within individual MCs were monitored with fluorescence and phase contrast microscopy (Figure 4C). The volume of media in the waste was measured at the end of the experiment to confirm that the DSCPM was successfully infusing at 1.5 μL/min for the entire duration.

**Figure 4.**
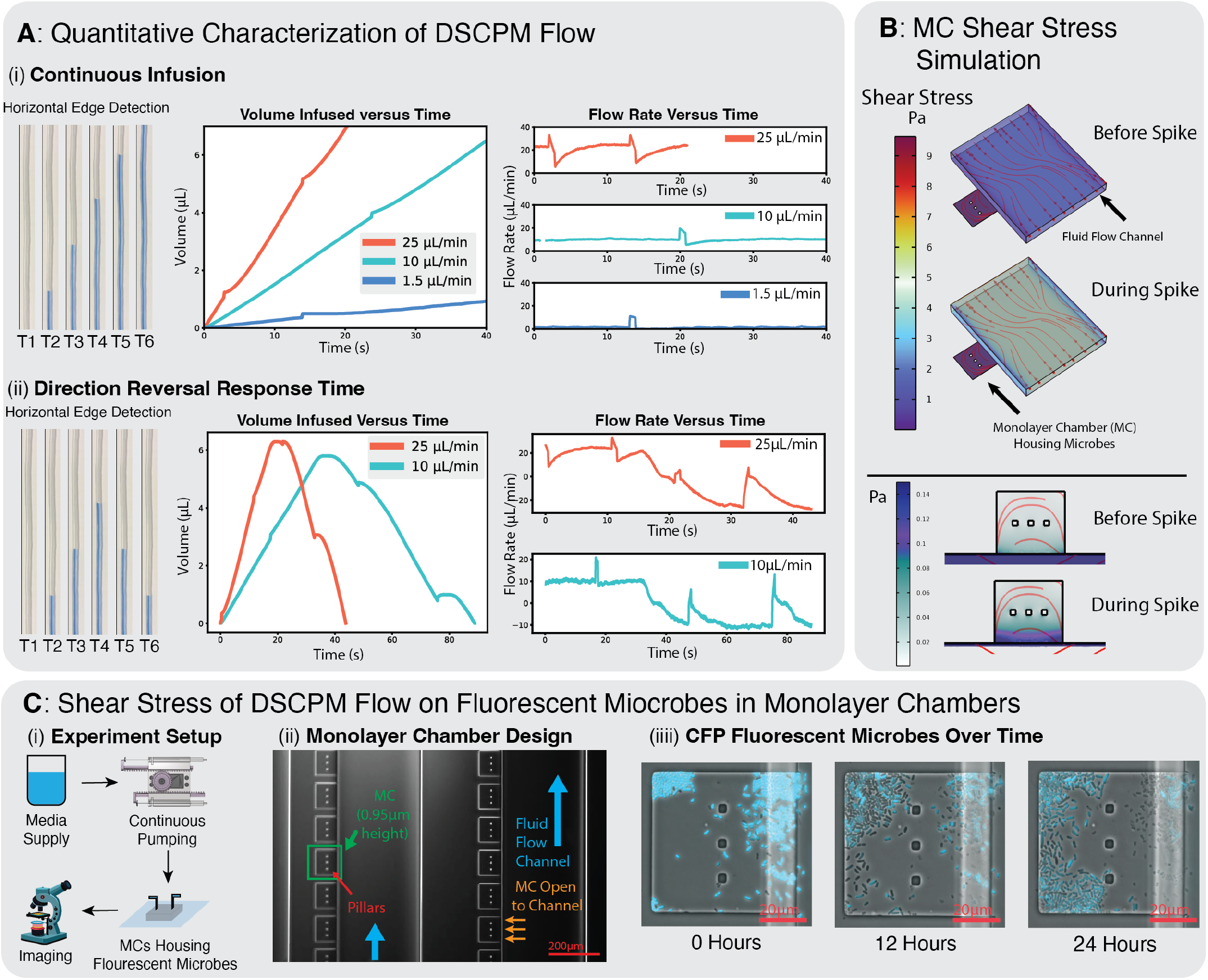
Characterization of DSCPM flow. A) Characterization of DSCPM Flow with direct flow rate measurements through image processing on video (60fps) of fluid in micro-bore tubing. i) Characterization of constant, forward flow for 25, 10, 1.5 μL/min. ii) Characterization of flow reversal response time for 25 and 10 μL/min B) Quantitative characterization of the shear stress experienced during one spike in flow rate by a cell trapping monolayer chamber (MC) with DSCPM flow via computational fluid dynamic (CFD) simulation. CFD simulation performed using COMSOL, assuming incompressible fluid. C) Proof of concept demonstrating the sustained maintenance of microbial consortium in photo-lithography microfluidic MCs for biological verification of DSCPM. i) Schematic of flow system setup for the tested use case of the DSCPM. ii) 10x phase contrast imaging of a MC containing PDMS microfluidic chip. iii) Images of one MC at three key time points throughout a 24-hour quorum sensing experiment which used microbes genetically engineered to constitutively express CFP. Images obtained by transposing fluorescence images onto corresponding phase contrast images (100x). The DSCPM infused media at 1.5μL/min for the entire duration.

Additionally, CFD (COMSOL) was used to model a spike in an MC to determine the range of shear stress a microbe would experience while being supplied media via the DSCPM. Shear stress in the channel ranged between 1 and 5 PSI. However, the shear stress in the MC itself stayed between 0.01 and 0.14 PSI, with the stress only ever exceeding 0.08 PSI on the very edge of the MC that faced the channel (See Figure 4B).

### Fluidic Multiplexing

Hardware support for perturbation automation can be integral to automating any fluidic system. Whether it is a single-strain consortium that requires a switch from plain media to media with a signaling molecule at a specific time or is a complex, multi-strain system which requires periodic perturbations between multiple medias, fluidic multiplexing (MUXing) with solenoid pinch valves can provide invaluable functionality to a flow system.

Three-way solenoid pinch valves are an excellent tool for building fluidic multiplexers because they can select between two inputs (Figure5). In theory, any N-to-1 MUX can be created with 2^*N*^ valves. Yet, the multiplexing configurations proposed and utilized in this study employ N - 1 valves to select between N sources. This is because the valves used for the DSCPM prototype (12V Three-Way Solenoid Pinch Valves from BEION/PreciGenome) cannot maintain the OFF state when pinching more than one cross-section of tubing. This is due to the physics of solenoids: they operate by generating a magnetic field when supplied with power that fixes the pusher in its extended ON state. When the solenoid is not provided with power, the pusher is not magnetically fixed to any position, and is held in its non-extended OFF state with a spring. With 12V BEION/PreciGenome solenoid valves, the activated magnetic field provides more than enough force to pinch multiple cross-sections of tubing when it is ON; however, the spring does not supply enough force to fully pinch more than one cross-section of (Shore A 50 durometer hardness) tubing in the ON state. If a BEION/PreciGenome valve were fitted with an additional external spring, different solenoids were used, or soft enough tubing was used, the 2^*N*^ valves rule would stand. Conceptually, the N-1 MUXes are analogous to one-hot encoding in logic design (See truth tables in Figure 5A), while binary encoding (2^*N*^) is usually structurally optimal.

**Figure 5.**
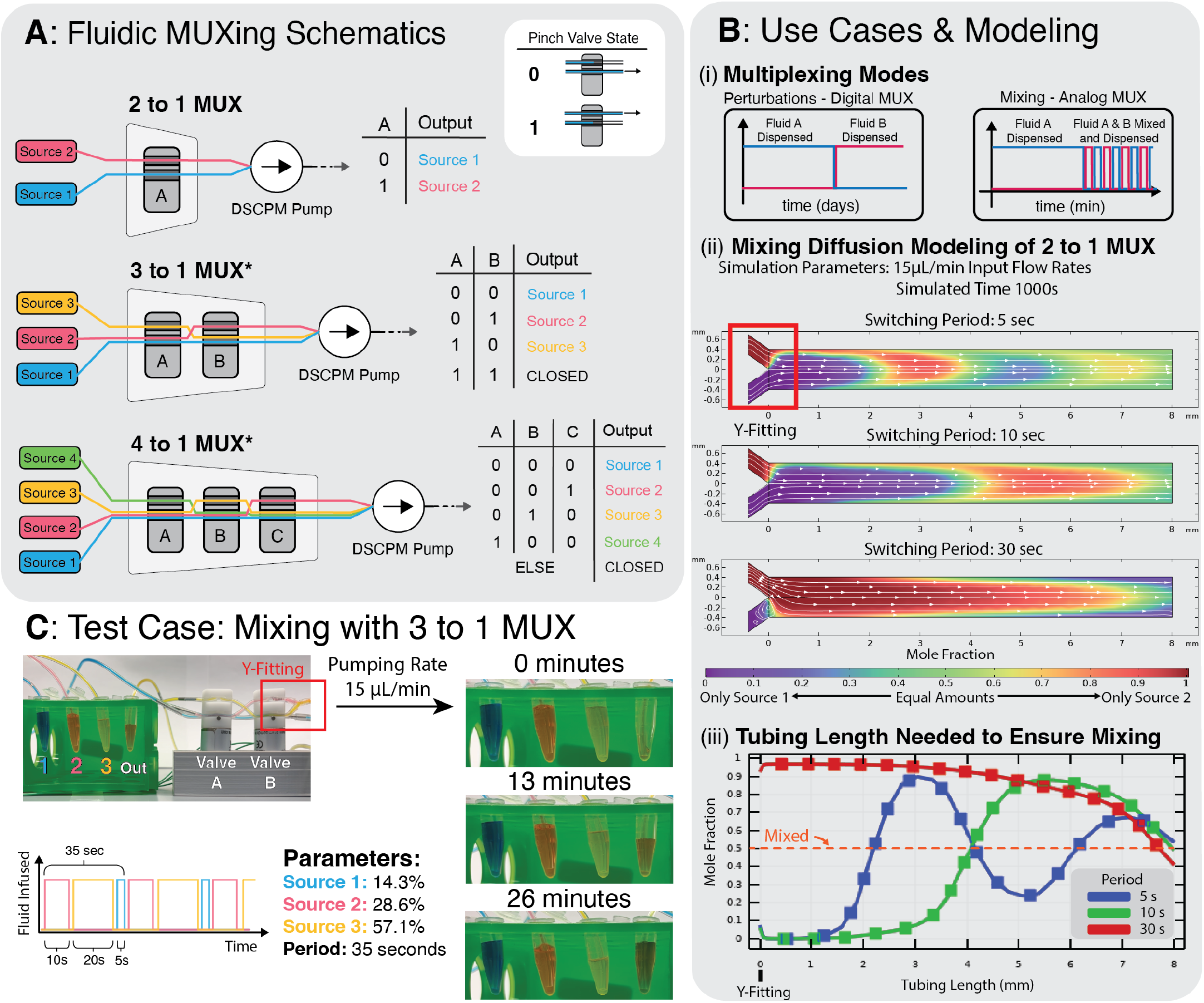
Fluidic Multiplexing with Three-Way Solenoid Pinch Valves. A) Fluidic multiplexing (MUXing) schematics with 3-Way Pinch Valves: “wiring diagrams” and valve-state / output truth tables. *The 3 to 1 and 4 to 1 MUX schematics pictured are the optimal obtainable schematics using BEION pinch valves and Shore A 50 durometer hardness tubing, due to the internal solenoid spring being unable to fully compress more than one cross-section of Shore A 50 tubing in the OFF state. These schematics follow the Select N+1 rule, while optimal MUXes using different valves or softer tubing could follow the Select 2 to the N rule. B) i) Main use cases for fluidic MUXes, which are perturbation between sources (“digital” MUXing), and mixing of sources (“analog” MUXing). ii) Mixing diffusion computational fluid dynamic modeling, which shows the diffusion of infused inputs in tubing with three different MUXing cycle periods. iii) Quantitative representation of the data from part (ii). The behavior of the mole fraction can be further modeled by over-damped oscillators, whose dampening coefficients can predict when convergence at a mole fraction of 0.5 is reached. C) Setup and results of an experiment using a 3 to 1 MUX, a DSCPM unit, and dyed input solutions. The DSCPM infused from the three reservoirs at 15μL/min, while the MUX “mixed” the three inputs with a cycle period of 35 seconds.

The fluidic MUXes allow for perturbations over time, which if applied periodically, can be implemented for mixing purposes (Figure 5B(i)). Biological applications for this include swapping the source media from one solution to another at a certain point or infusing a ratio between multiple medias. COMSOL simulations were performed to quantify diffusion over time between mixed sources as they were infused. Based on the MUXing cycle period, the fluid mole fraction (of source A to source B) in the tubing was observed to oscillate along the tubing position (Figure 5B(ii)). The results showed that diffusion occurs more rapidly for shorter periods (T=5 seconds).

As a proof of concept, a 3 to 1 fluidic MUX was constructed and programmed to multiplex between three sources in a 2:4:1 ratio, dyed red, yellow, and blue respectively. The MUX was placed in series with the sources and the DSCPM, which infused for the duration of the experiment at 15 μL/min. The MUXing cycle period programmed was 35 seconds long. Imagery of the three source reservoirs and the target reservoir was captured intermittently throughout the experiment to confirm that each reservoir’s expected volume was infused.

## Discussion

This study successfully designed and prototyped an open-source pumping system capable of providing continuous precision flow and multiplexing between sources. The flow generated by the DSCPM was quantitatively characterized and tested in a real-world application involving microbes housed in cell trapping MCs.

The DSCPM addresses the need for a continuous and precise pumping solution while costing significantly less than pumps that achieve similar flow rates. The easily integrable fluidic multiplexers further enhance the advantages of the DSCPM system and reduce the need for additional mixing and perturbation components. Dual-syringe rectifying pumps have been done before (See New Era Pump Systems DUAL-NE-1000X Continuous Infusion Syringe Pump System); however, each market available solution that employs this concept uses check valves for flow rectification, which disrupt flow laminarity at low flow rates (<100*μ*L/min). Additionally, these solutions require two independent motor systems, given that their syringes do not operate as direct inverses as one another. With larger syringes, switching direction has a delay, likely due to the elasticity of the syringe itself. To fix this, these systems use a mechanism called syringe priming, which means that the refilling syringe refills faster than the infusing syringe infuses, so that it has time to prime itself for infusion before it switches. This results in a less optimal system. These solutions generally consist of a “kit” of rectifying equipment meant to be installed to traditional syringe pumps; meaning that one continuous infusion unit will costs thousands of dollars, due to the market value of traditional syringe pumps and the fact that two pumps are required.

Determining what experimentally constitutes sufficiently laminar flow generation in low-volume and low-flow fluidics remains somewhat unresolved in the field due to the lack of precise sensors in the low-flow rate regime. Traditional fluid mechanics ascertains that flow is laminar if the Reynolds number is between 0 and 2320; however, experimentally collecting data that can yield Reynolds numbers for low flow rates is not always feasible. However, as shown above, a golden standard product’s specifications suggest that a threshold of one minimum pump increment every 30 seconds can be considered adequately laminar. It is crucial to note that this is an arbitrary threshold, and flow generation, regardless of precision, will always be discretized into increments.

Using our flow rate versus time data and computational fluid dynamics to analyze the shear stress exhibited on cell trapping MCs, we showed that during infusion, the shear stress is below 4^*−*2^ Pa, and during the valve state switch induced flow rate spike, the maximal shear stress (on the MC and channel interface) increases to around 12^*−*2^ Pa. Literature shows that some shear sensitive mammalian cell types, such as embryonic stem cells and primary neurons, are detrimentally affected by any shear stress exceeding 10^*−*4^ Pa, with hepatocytes also being negatively affected by stress greater than 10^*−*2^ Pa^29^. While the shear stress experienced by the MCs in our microfluidics due to the flow rate spikes of the DSCPM may make the shear conditions unsuitable for specific cell types, this is primarily due to the geometry of our microfluidics, given that they were designed to house bacteria, which are much more resilient than mammalian cells.

CFD simulations yielded a shear rate on surfaces within the PDMS microfluidic: a maximum of <500 *s*^*−*1^ in the MC. A previous study that measured what shear rates would cause bacterial detachment from a flat glass wall measured shear rates on the order of magnitude of <10,000 1/s^30^, so it can be inferred that microbes in a PDMS cell trapping MC would be unlikely to be disturbed. Our proof of concept experiment, which consisted of CFP microbes in MCs, shows that in spite of the flow rate spike that is induced by the free flow preventing two-way valve, the DSCPM flow is adequately steady, or “laminar”, to maintain proliferation of the microbes. Fluorescence is observed at all time points during the experiment, and the microbe population increases from the beginning to the end.

The DSCPM hardware utilizes a 3D-printed linear actuation mechanism to convert the rotational motion from a motor into linear motion. The design was adapted from an existing open-source Linear Servo Actuator. The DSCPM prototype uses a GoBilda High Torque Servo Motor, a DC motor with built-in rotary encoders, controllable by an Arduino library using position (usually angle) as an argument. Typically, servo motors turn one degree per increment in the code; however, GoBilda High Torque Servo motors turn 1.5 degrees per increment, making the minimum increment 1.5 degrees. The DSCPM designs that prevent free flow (DSCPM 2-3/1-2 and DSCPM 2-2) introduce periodic pressure spikes due to the added free-flow prevention fluidic logic. Future iterations of the free-flow-preventing DSCPM design could minimize these spikes by reducing the inner diameter of the tubing since less volume would be displaced. This could involve designing custom solenoid pinch valves for smaller tubing sizes or creating new logical schematics that are equally functional.

Future iterations of the DSCPM that utilize a motor with a smaller angular minimum increment could significantly increase flow generation precision, and enable employment of of larger volume syringes while keeping the same achievable flow rate range. The commonly used NEMA-17 stepper motor (200 steps/revolution with 1/16th microstepping) with a driver capable of supporting micro-stepping can reach 3,200 steps per revolution. With this motor, DSCPM is calculated to achieve the resolution of expensive 0.5mL syringes with disposable 1mL syringes. Therefore, the following versions of the DSCPM will benefit from higher-resolution motors that enable a more comprehensive range of applications. While the use of larger syringes could introduce a need for syringe “priming” at the state switch, which would be detrimental to the mechanism, this design iteration would be worth trying; additionally, improving the precision of the linear actuation mechanism by decreasing the rotational motion to linear motion ratio could achieve greater precision with the same motor setup.

## Methods

### DSCPM Design and Fabrication

#### Electronics and 3D Printed Parts

The DSCPM housing STL file was sliced (prepared for printing) with Ultimaker CURA and 3D printed with acrylonitrile butadiene styrene (ABS) filament on a Creality Ender 3 Printer. The base was printed on its back to ensure no supports were needed in the pusher grooves. The pushers were printed with the teeth facing up, and the gear was printed on its back (orthogonal to its rotational axis). The bottom and sides of the pushers were sanded with a Dremel tool until they could slide into and out of their grooves smoothly. The provided servo gear horn was cut (using wire cutters) just above the innermost hole on each side. A large hole was drilled into the center of the gear, and two smaller holes were drilled in line with the gear horn screw holes. The gear horn was placed onto the gear (so that the base fit into the large hole in the gear), and two #6 screws were used to fasten the gear to the provided servo gear horn. The circuit was assembled on a breadboard according to the schematic shown in (Figure S1).

#### Initial Hardware Assembly

The DSCPM code was uploaded to the Arduino micro-controller with the EEPROM initializers (lines 16-18) uncommented, and the EEPROM statements (lines 29-31) below them commented. The serial command “123” was passed (to “start” the pumps), and then the serial command “0” was passed (to “pause” the pumps, which stores the motor position in the EEPROM of the Arduino). Then, the code was re-uploaded with the EEPROM initializers commented and the EEPROM statements below uncommented. If any syringe that is not 10μL is used, the variable that stores the syringe inner diameter (“innerdiameter”) on line 55 should be changed accordingly. If an alternative servo is used, the instructions in the comments from lines 64 to 68 should be followed, and the variable “degper180” on line 70 should be changed accordingly.

The gear was fitted onto the servo, and the serial command “123” was given to start the pumps. When the position reached its user-designated minimum (default position is 5), the pumps were paused with the serial command “0”. The gear was removed from the servo, the first syringe was placed, and the first pusher was slid into its groove. The gear was fitted back onto the servo, with the first pusher’s teeth aligning with it at the minimum position of the syringe’s plunger. The pumps were un-paused by passing any integer below 25 (this sets the pumps’ flow rate). They were then re-paused at their user-designated maximum position (the default is 70). The process of adding the first syringe and plunger was repeated for the second syringe. This process ensures that the placement of the pushers will give the syringes their optimal range of motion as the pump cycles. This process should be repeated if the maximum or minimum angle parameters are ever altered. In the event of any updates, the most recent software and accompanying instructions will be in the DSCPM repository of the Sam Oliveira Lab GitHub (https://github.com/SamOliveiraLab/dual-syringe-pumps).

The electronics were assembled based on the circuit schematic in Figure S1. The solenoid pinch valves are fitted with flyback diodes and an overdrive function (implemented in the code) to keep their temperature cool (Figure S3).

#### Initial Fluidics Assembly

If using 10 μL syringes or smaller, fill each syringe with water. The volume of one syringe infusion is smaller than that of the Luer tip chamber fitted onto the syringe, meaning that the fluid being pumped will never actually reach the syringes (work done will be by suction). Syringes of 10 μL and smaller are extremely delicate due to the small inner diameter, and moving the plunger when the syringe is dry can damage it, so having water present is essential. Next, fit the Luer tips onto the syringes (it is a friction fit). It is recommended that parafilm be applied outside of the junctions to ensure that they are airtight. Then connect each Luer tip to a Y-connector using 1/8” OD, 1/16” ID tubing.

This next part describes how to assemble a DSCPM 2-3/1-2 configuration using 1/8” OD rectifying (3-way) valves and a 1/16” OD free flow prevention valve: Connect one outlet from each Y-connector to a third Y-connector using the same tubing. Repeat the process with the remaining outlets on each syringe’s Y-connector. One will connect to the source and the other to the system. Connect one Y-connector with the remaining outlet to the source. Using a 1/16” ID to 1/32” ID tubing converter (our method is outlined in Figure S4), connect the remaining unused Y-connector outlet to the 1/32” tubing provided with the two-way solenoid pinch valve.

### DSCPM Setup

#### Subsequent Fluidics Assembly and Sterilization

After the first time the fluidics are assembled, they should not be taken apart for sterilization purposes. The tubing was sterilized by removing the valves, and 10% bleach was infused through the pump system, making sure to manually pinch tubing in each branch closed periodically to ensure flow through the complimentary branch. The reason for this is that if one branch has greater resistance than the other, there may be inadequate sterilizing of the higher resistance branch due to a lower flow rate. The system was then rinsed with deionized water three times following the same procedure as the bleach infusion. This sterilization procedure was observed to be adequate to prevent contamination.

### Monolayer Chamber-Containing Microfluidic Chip Design and Fabrication

The Polydimethylsiloxane (PDMS) based microfluidics were fabricated by designing a two-layered device to house the channels and chambers. A CAD software tool (KLayout) was used to design the two layers (Figure S5). Each design was written on a Soda Lime glass mask using a mask writer (DWL66 by Heidelberg). SU8-2002 was mixed with pure 99+% cyclopentanone in a 4:6 ratio then spin coated onto an Silicon wafer after plasma treating the wafer with O_2_ (Plasma Asher M4L by PVA TePla America), to achieve a thickness of 0.95 μm resist that was used for patterning of the chambers layer. After that, the wafer is soft baked at 65°C/95°C for 30 seconds and 3 minutes respectively. The wafer was placed under its designed mask in a mask aligner (MA6 by Karl Suss). The wafer was developed for 90 seconds after post exposure baking at 65°C/95°C for 90 seconds and 120 seconds respectively. The wafer was Plasma treated again in order to prepare the wafer for the second layer. Spin coating was done on the wafer using SU8-2025, to achieve a channel layer thickness of 25 μm. The wafer was soft baked at 65°C/95°C for 3 minutes and 10 minutes respectively. The wafer was then placed in the mask aligner under the channels mask and aligned properly. Post exposure baking was done at 65°C/95°C for 90 seconds and 4.5 minutes respectively. Then the final developing step was done for 4.5 minutes followed by hard baking at 200°C for 1 hour. PDMS was later mixed in a 10:1 ratio of the base to the curing agent then poured onto the wafer and placed in a 75°C oven for 12 hours. This led to the negatives of the wafers printing onto the PDMS chips, which were later diced and plasma treated to bond onto glass slides.

### Experimental Methods

#### Quantitative Flow Characterization with Micro-Bore Tube Infusion and Video Processing

The DSCPM was configured to infuse into Tygon Non-DEHP Micro Bore Tubing (0.030” ID), which was secured against a white background. Water with blue food coloring was infused through the pumping system, and a 150mm long section of micro bore tubing was filmed with a smartphone. The videos were processed using an algorithm that analyzed each frame with a vertical edge detection kernel, meaning that fluid position along the length of tubing over time was quantified. Then, using the inner diameter of the tubing, volume infused over time data was calculated from the fluid position versus time data. Additionally, the derivative with respect to time of the volume infused versus time data was taken to calculate the flow rate over time. During data collection, the pumps were controlled with serial connection from a computer.

#### Quantitative Flow Characterization with Computational Fluid Dynamics

Through the application of Computational Fluid Dynamic (CFD) modeling, the laminarity and shear stress of a proposed pumping mechanism, along with its associated flow rectification method using pinch valves, were thoroughly examined. The system’s flow rate measurements, obtained via video processing, were modeled in COMSOL Multiphysics (V6.1 with microfluidics module). The tubing was modeled in COMSOL under the 3D laminar flow interface to investigate the potential disruption of the laminar flow profile by flow rate spikes induced by the pinch valves. The input, a time-dependent fully developed volumetric flow rate, was derived from the collected image processing data. The system was assumed to be incompressible, and the Reynolds-Averaged Navier-Stokes (RANS) turbulence model was utilized. The average Reynolds numbers were reported for three examined flow rates at a point on the centerline.

The impact of flow rate spikes within the microfluidic cell trapping MCs and their potential effect on housed microbes were assessed by modeling the MCs in COMSOL. GDS files for each device layer, used for photolithography manufacturing, were processed in Autodesk’s Fusion 360 to create a .stl of the MC. A Multiphysics model of the MC was established using the 3D laminar flow interface and transport of dilute species. The input flow rate, derived from image processing, and species transport concentration of 0.5M were set under incompressible flow with RANS turbulence modeling. The model yielded shear rates and stress values on the 3D model surface.

#### Quantitative Flow Characterization with Series Pressure Sensing

The Honeywell ABP2 Series Differential Pressure Sensor (ABP2DDAN001PD2A3XX) was connected to an Arduino Nano micro-controller (Figure S2). The sensor was set up in a monometer configuration in series with the DSCPM unit and the target, which was just an additional length of tubing. The pressure (percent of sensor maximum, not calibrated to any unit) was collected at a 10 ms interval, using an Arduino Uno and an adaptation of the code found in the sensor specification sheet (www.mouser.com/datasheet/2/187/Honeywell_01292021_ABP2_Series_Datasheet_Issue_C-1991028.pdf).

The differential pressure sensor’s relative pressure against its maximum capability is ± 1 psi. The sensor was integrated in series with the system in a monometer configuration, meaning that the first input was connected in series with the DSCPM, and the second input was linked to a closed air reservoir at 1 atm (14.7 psi). An equation was derived using the sensor specifications to convert percent pressure from maximum (the raw output), wherein equal pressures yield 50%, and a one PSI difference corresponds to 100%. The equation yields the absolute pressure in series with the DSCPM.

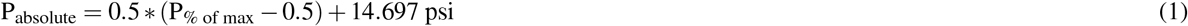

In the original data, larger frequency components were observed due to slow changing, overarching pressure gradients in the pressure sensing setup. The data was corrected for main noise and low frequency oscillations in the measurement through frequency domain analysis. Filtering out the larger frequency components was necessary to perform analysis on the data which uses the standard deviation of relative pressure as a rudimentary quantification of laminarity for comparison between the HA PhD 2000 and the DSCPM. The standard deviation of relative pressure for the HA PhD 2000 flow was expected to remain relatively similar regardless of flow rate. The standard deviation of the DSCPM flow was expected to increase with flow rate due to the increase in frequency of valve state switch-induced “pulses”. Furthermore, since the pulses are the main disruption of laminarity in the flow produced by the DSCPM, peak detection was performed on the noise-corrected relative pressure data, and average peak height and standard deviation were calculated for each of the three sampled flow rates. Theoretically, the pulse magnitude should be the same across all pulses at all flow rates, since the same amount of fluid is being displaced at the same rate (the two-way solenoid pinch valve in the DSCPM 2-3/1-2 schematic takes the same amount of time to close and open regardless of the flow rate).

#### Maintenance of Microbial Consortia with the DSCPM

Bacteria cells (*Escherichia coli strain*) that were genetically engineered (plasmid obtained from addgene, and cells transformed by the DAMP Lab) to express Cyan Fluorescent Protein (CFP) constitutively were infused in media into the main channel of a microfluidic that contains cell trapping MCs. The chip was then centrifuged with the MCs facing outwards from the center of the centrifuge, which forced the microbes into the MCs. A DSCPM unit was sterilized and set up to infuse from a media-filled reservoir into the microfluidic chip. Media was infused through the chip at 1.5 μL/min for 24 hours as the chip was placed in an inverted microscope (Nikon Eclipse Ti2) for imaging using a software application (NIS Elements, Version 5.30.00) to capture the MCs layer by taking images under phase contrast and CFP every 5 minutes and 3 minutes respectively. For these chips specifically, this flow rate is on the low end of guaranteeing adequate delivery of nutrients to the microbes while avoiding detrimental shear stress on the MCs.

#### Flow Behavior Characterization with the 3 to 1 MUX

Water from three distinct reservoirs, dyed blue (Source 1), red (Source 2), and yellow (Source 3), was infused through the DSCPM/MUX system until the volume of the system had been completely infused with fluid. The DSCPM was paused, and the source and output vessels were exchanged for 0.4 mL micro-tubes to allow increased visibility of small volume changes. The system was then un-paused and filmed for approximately 30 minutes with a DSCPM flow rate of 15 μL/min, and a MUXing cycle which infused a 1:2:4 mix between sources 1 (blue), 2 (red) and 3 (yellow) respectively. The MUXing cycle had a period of 35 seconds, making source one, two, and three’s respective periodic infusion durations 5 seconds, 10 seconds, and 20 seconds.

#### Multiplexer Mixing Diffusion Modeling with Computational Fluid Dynamics

A 2 to 1 MUX was modeled using COMSOL Multiphysics (v6.1) to achieve a 1:1 mix of two dilute species inputs. The dimensions of the Y-fitting and 8mm of 1/32” ID tubing were represented as a 2D model, incorporating 2D laminar flow and transport of dilute species in a time-dependent study (t=1000 seconds). The inlets were configured as a square wave with a 15μL/min baseline, amplitude, and periods of 5, 10, and 30 seconds. The two inputs had equal flow rates but were 180 degrees out of phase. The species concentrations were set at 0.50M, with initial boundary concentrations set to zero.

## Acknowledgements

The authors gratefully acknowledge funding support from the National Science Foundation NSF (#4500004978), the Defense Advanced Research Project Agency DARPA (#D24AP00330-00), the iGEM Impact Grant Award (2023), and the Department of Defense DoD STEM FY20 Award (HQ00342110008). The authors would also like to thank the Biological Design Center of Boston University, STEM Pathways of Boston University, and the following BostonU-HW 2023 iGEM members: Margherita Piana, Arjavi Vyas, Olivia Gibson, Daniel Oh, Yazmin Camacho, Diana Arguijo, and Juan José Robayo Yepes for discussions within the iGEM project where this work originated. Finally, we thank Prof. Douglas Densmore (CIDAR and DAMP Lab PI) for the technical and funding support for the iGEM project where this work originated.

## Author Contributions Statement

S.M.D.O. conceived the primary use case and desired hardware metrics for the DSCPM. K.A.W. designed and fabricated the pump and multiplexer hardware, and wrote the pump and multiplexer software. K.A.W. conceived and performed the flow rate sensing experiments with visual tracking, and the multiplexing experiment. K.A.W. and Y.H.P. conceived and performed data analysis for the pressure sensing experiments. K.A.W conducted the pressure sensing experiments. Y.H.P performed all computational fluid dynamic simulations. W.A. designed and fabricated the microfluidics with the cell trapping MCs. W.A collected microscope images while performing the biological verification experiment. H.L.G. and S.M.D.O. acquired funding and lab space. K.A.W., Y.H.P., and W.A. contributed to writing the manuscript. All authors edited the manuscript.

## Additional information

S.M.D.O. co-founded Doroth, an AgTech start-up focused on developing DNA-based sensors and automation technologies to monitor biological targets in crop fields remotely. Y.H.P. is an employee of BioSens8, a start-up focused on wearable microfluidic sensing technologies. The remaining authors declare no competing interests.

While preparing this work, the authors used Generative AI (OpenAI’s Chat GPT and Microsoft’s Copilot) to improve the readability of the material. They also used generative AI to aid in writing the image processing python script for the flow rate versus time results. After using these tools/services, the authors reviewed and edited the content as needed and take full responsibility for the content of the published article.

## Notes

https://github.com/SamOliveiraLab/DSCPM

