## Supplementary Information for "A Continuous, Low-Flow, and Multiplexing Pumping System for Microfluidics Applications"

**Affiliation:**
[1]Department of Biomedical Engineering, Boston University, Boston, MA, 02215, USA
[2]Department of Electrical and Computer Engineering, Boston University, Boston, MA, 02215, USA
[3]Joint School of Nanoscience and Nanoengineering, North Carolina A&T State University, Greensboro, NC, 27401, USA

### Table of Contents:

**Supplementary Methods:**
1. Derivation of Steady/Laminar Flow Metric with Harvard Apparatus

**Supplementary Figures:**
1. DSCPM Circuit Schematic
2. Pressure Sensing Experiment Data, Circuit, and Setup
3. Overdrive Function and Flyback Diode for Solenoid Pinch Valves
4. Fluidics Assembly: Adapting from Y-connector (fits 1/16” ID tubing) to 1/32” ID tubing
5. PDMS, MC containing Microfluidic Chip CAD and Fabrication Details
6. DSCPM CAD Images

**Supplementary Tables:**
1. Bill of Materials
2. Reachable Steady/ “Laminar” Flow Rates by the DSCPM

**Appendix A: STL Files**

**Appendix B: Arduino Codes**

### Supplementary Methods

**Supplementary Section S1:**

The specifications of the PhD 2000 cite the metric of 0.18 µm per minute as the minimum pusher travel rate. Equation (1)

From this, we can calculate:

$$\frac{1\ \mu{-}step}{0.08\ \mu m\ linear\ motion} * \frac{0.10\ \mu m\ linear\ motion}{1\ minute} = 2.19\frac{minimum\ increments}{minute} \quad (1)$$

This number seemed constant regardless of the volume infused per minimum increment. Given this, this study's accepted metric for sufficiently laminar flow is a maximum interval between minimum increments of 27.34 seconds.

The 3D-printed linear actuator from which the DSCPM housing was adapted (www.thingiverse.com/thing:3170748) yields 0.256 mm of linear motion per degree of motor rotation. The high torque servo motors used in the DSCPM prototype had a minimum increment of 1.5 degrees. Given this, the minimum laminar flow rates for the configurations listed in Table S1 were calculated by:

$$\pi * \left(\frac{Syringe\ ID}{2}\right)^2 * \frac{0.256\ mm}{degree} * \frac{0.15\ degrees}{minimum\ increment} * \frac{1\ \mu L}{mm^3} * \frac{2.19\ minimum\ increments}{minute} \quad (2)$$

$$= minimum\ flow\ rate$$

The DSCPM housing can generally hold syringes from 0.5µL to 1mL since the standard dimensions for those syringes all have the same outer diameter. This means that the DSCPM can be customized based on the required flow rate range for an application, without additional altercations to the housing.

### Supplementary Figures

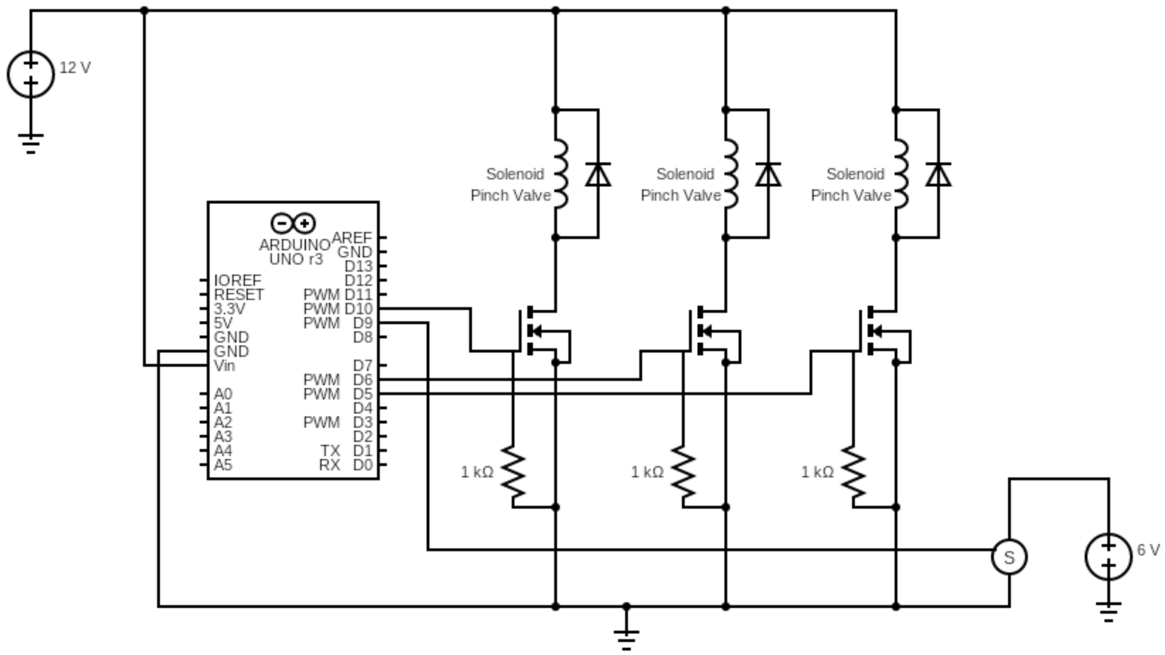


**Figure S1:** DSCPM 2-3/1-2 Circuit Schematic. All diodes are IN4007, and the Servo Motor is a 6V High-Torque Servo Motor. Two amp power supplies were utilized. When making a DSCPM 2-2, the left-most solenoid (connected to Pin D10) is omitted.

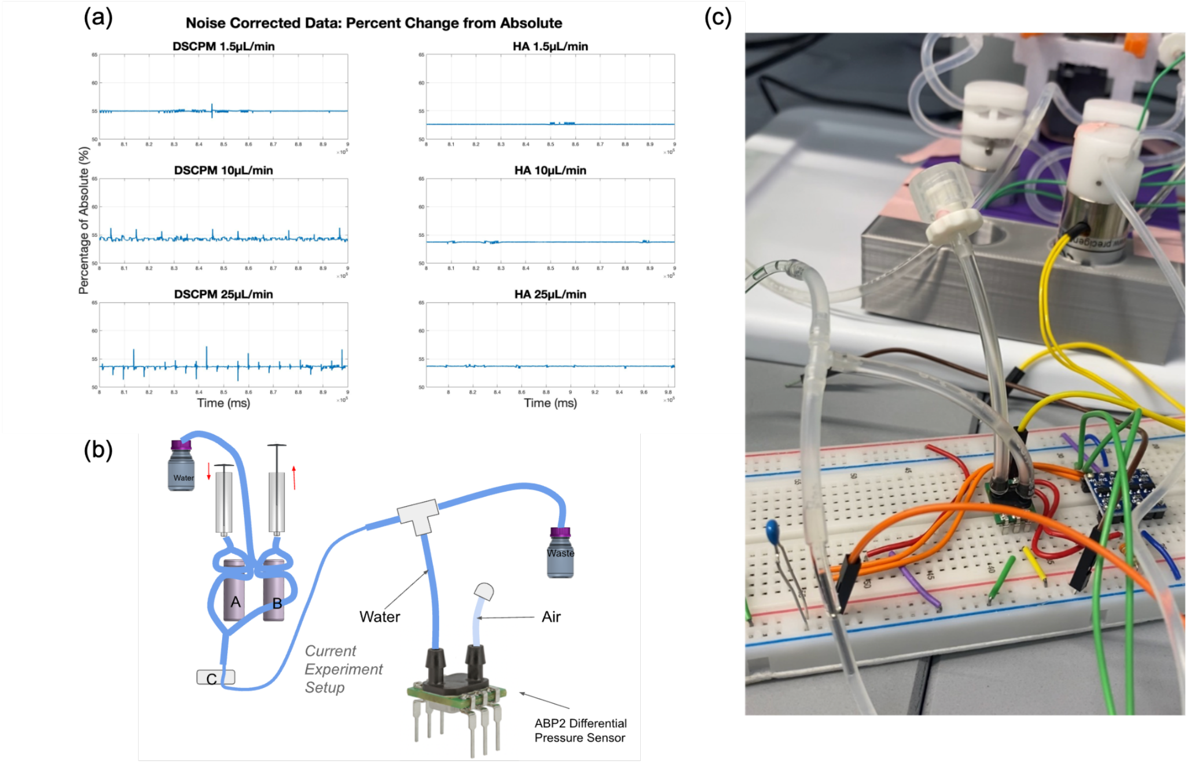


**Figure S2:** Pressure Sensing Experimental Data and Setup. (a) Noise-corrected pressure vs. time data for the DSCPM and the HA PhD 2000 fitted with a 60mL syringe. (b) Fluidics schematic of the Honeywell ABP2 Series Pressure Sensor in a monometer configuration in series with the DSCPM. (c) Photo of the experimental setup.

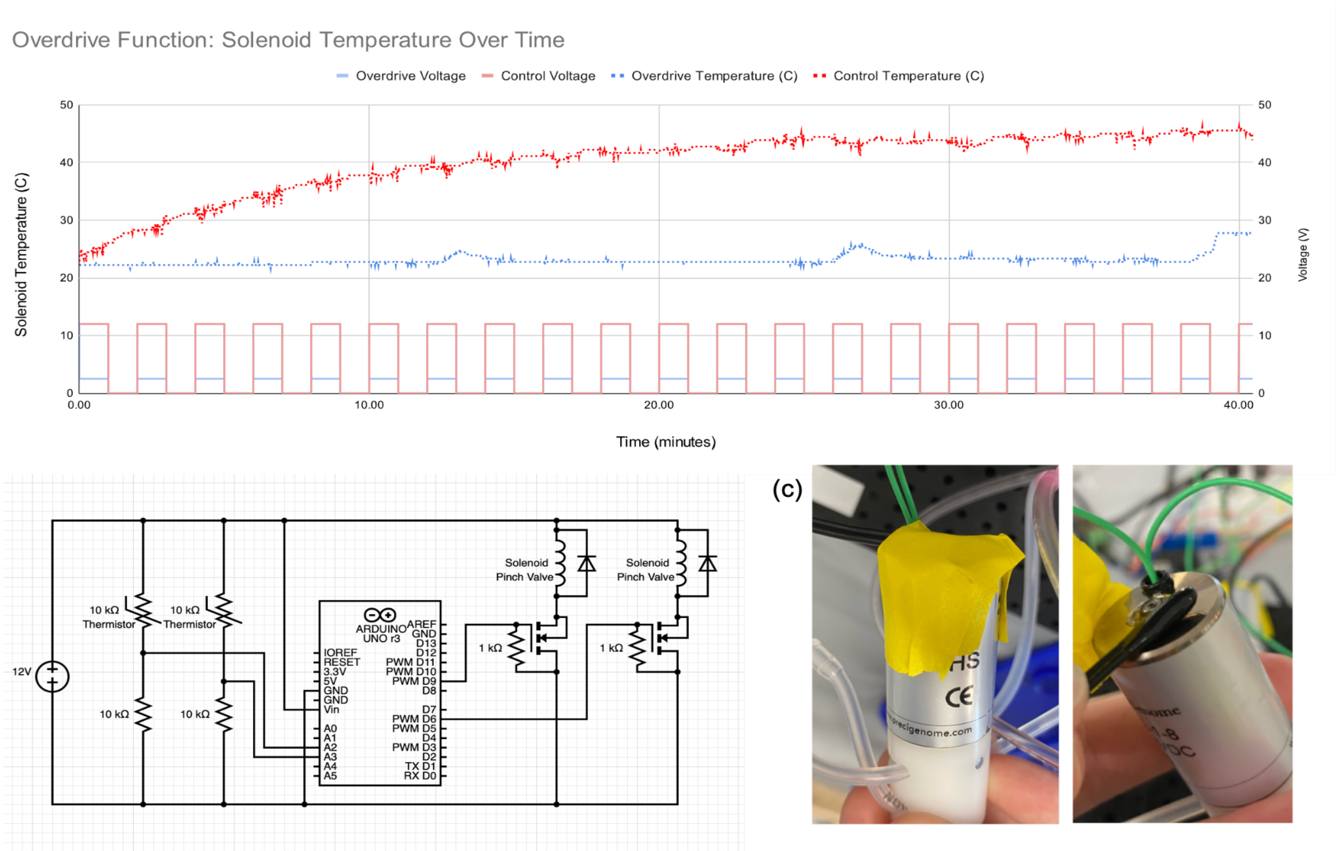


**Figure S3:** Overdrive Function for long-term temperature control of BEION/PreciGenome 12V Solenoid Pinch Valves. Measurements were taken with a 10K thermistor. (a) Solenoid Temperature versus time transposed with voltage duty cycle versus time. (b) Temperature Measurement setup circuit. (c) Thermistor placement on the base of the solenoid to maximize surface area in contact with the thermistor.

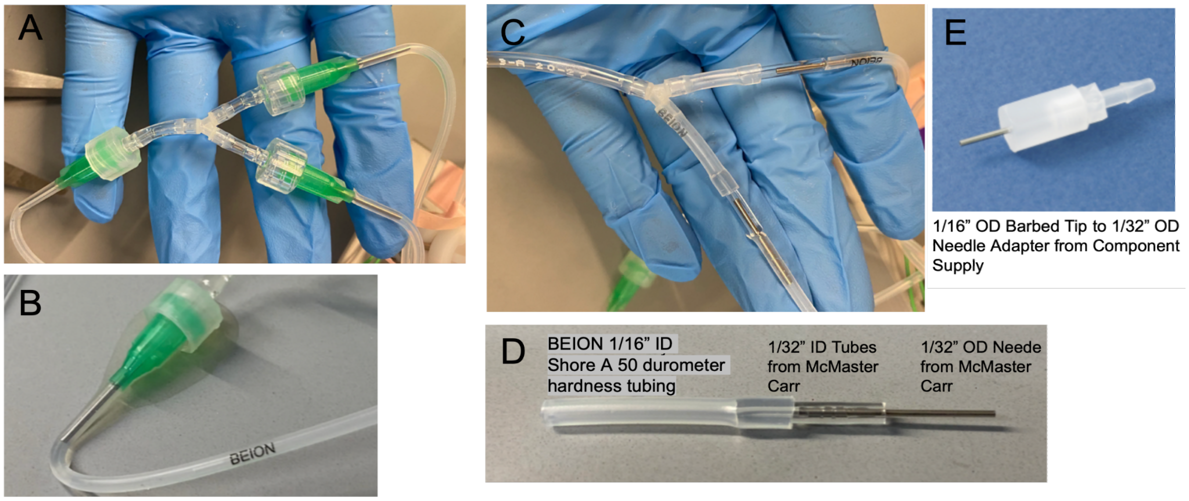


**Figure S4:** (a) Adapting from a Y-connector (meant for 1/16” ID tubing) to 1/32” ID tubing with multiple Luer connections, which have a lot of dead volumes, which can introduce the risk of residual bubbles in the system from the assembly. (b) Leakage of media from a Luer connection. (c) Adapting from a Y-connector to 1/32” ID tubing with custom adapters. (d) The custom adapter schematic. The needle was obtained from a Luer tip/hub to needle adapter by compressing the plastic with pliers until it cracked and pulled the needle out. (e) A market-available adapter from 1/16” ID to 1/32” ID has no dead volume but is quite expensive compared to the custom adapters we made.

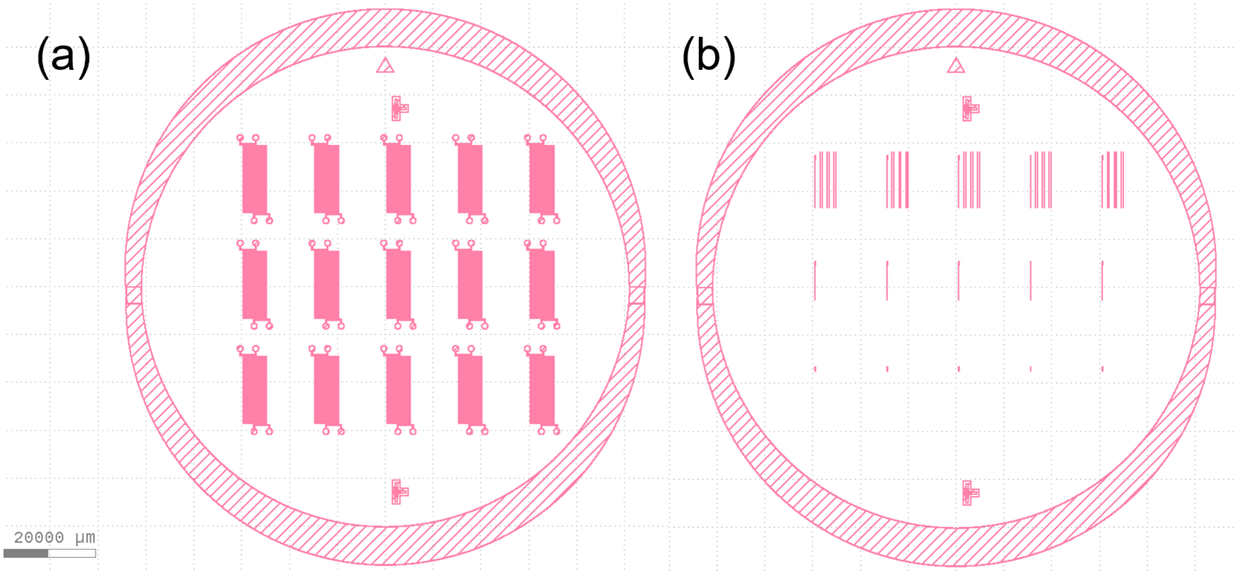


**Figure S5:** (a) CAD model of 15 chips used to imprint on a photomask using the Mask Writer. This layer acts as the second layer of the PDMS chips, the channels. (b) The first layer of the PDMS chips with different numbers of MCs per row.

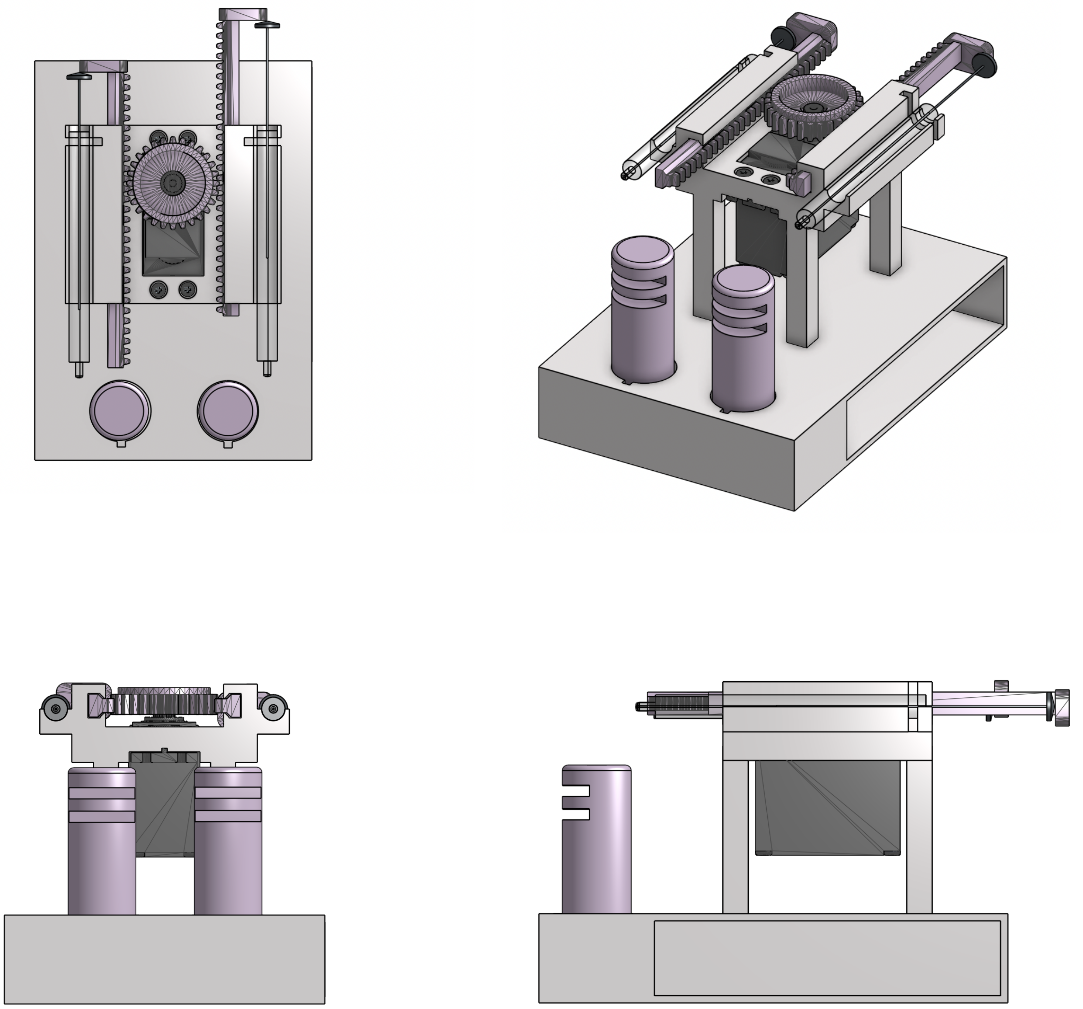

**Figure S6:** Top, Isometric, Front, and Left views of a DSCPM CAD drawing.

### Supplementary Tables

| **Bill of Materials** | | **KEY:** | **Both Configurations**<br>**DSCPM 2-3/1-2 Parts**<br>**DSCPM 2-2 Parts**<br>**N to 1 MUX** |
|---|---|---|---|
| **Part** | **Vendor** | **Amount** | **Price per Unit** |
| **Electronics Components:** | | | |
| Arduino Uno | Any | **1, 1** | ~ $30 |
| IN4007 Diodes | Any | **3, 2, (N - 1)** | < $1 |
| 1 kΩ Resistors | Any | **3, 2, (N - 1)** | < $1 |
| N-Channel MOSFETS | Any | **3, 2, (N - 1)** | < $1 |
| 3-Way Solenoid Pinch Valve | PreciGenome*<br>BEION | **2, (N - 1)** | $118* (PreciGenome)<br>~ $30 (BEION) |
| 2-Way Solenoid Pinch Valve | PreciGenome*<br>BEION | **1** | $108* (PreciGenome)<br>~ $30 (BEION) |
| High Torque Servo Motor | GoBilda | **1** | ~ $30 |
| 6V, 2A AC/DC Power Supply | Any | **1** | ~ $12 |
| 12V, 2A AC/DC Power Supply | Any | **1** | ~ $12 |
| **Fluidics Components:** | | | |
| 10 μL Syringe (Model 701 LT) | Hamilton | **2** | $36 |
| Y-Connectors for 1/16" ID Tubing | Cole Parmer | **4, (N - 1)** | < $1 |
| 1/16" ID Silicone Tubing | Any | **n/a** | < $1 |
| 1/16" ID Silicone Tubing: Shore A 50 Durometer | Any | **n/a** (comes with PreciGenome valves) | < $1 |
| 1/32" ID Silicone Tubing: Shore A 50 Durometer | Any | **n/a** (comes with PreciGenome valves) | < $1 |
| Luer Tip/Hub - 1/16" Barbed Tip Adapter | Any | **2** | ~ $2 |
| Luer Tip/Hub - 21 Gauge Needle Adapter | Any | **4** | < $1 |
| **Total Prices:** | **DSCPM 2-3/1-2:**<br>**DSCPM 2-2:**<br>**N to 1 MUX:** | **~ $250**<br>**~ $220**<br>**~ $[30(N - 1) + 45]**<br><br>**^ Optimal Prices** | **~ $500***<br>**~ $400***<br>**~ $[118(N - 1) + 45]***<br><br>**^ With PreciGenome Valves*** |

**Table S1:** Bill of Materials. *The DSCPM prototype uses BEION syringes purchased from PreciGenome, which cost about $100; however, the same valves are available from other suppliers for cheaper.

| **Syringe Volume (µL)** | **Syringe ID (mm)*** | **Volume per Minimum Increment (µL)** | **Minimum Flow Rate (µL/min)** |
|---|---|---|---|
| 0.5 | 0.10 | 3.0E-3 | 7.0E-3 |
| 1.0 | 0.15 | 5.0E-3 | 14.0E-3 |
| 10 | 0.49 | 7.1E-3 | 152.0E-3 |
| 1E3 | 4.78 | 6.89 | 15.12 |
| 5E3 | 12.07 | 43.94 | 96.41 |
| 10E3 | 14.50 | 65.41 | 139.14 |

**Table S2:** Reachable Laminar Flow Rates by the DSCPM with various market-available syringe sizes. * Each inner diameter corresponds with the standard dimensions for each syringe size. The minimum Laminar Flow Rate depends on the inner diameter, not the syringe volume.

### Appendix A: STL Files

In appendix A: The .STL files for 3D FDM printing the housing for the DSCPM along with .GDS files needed to create the microfluidic chambers using cleanroom photolithographic techniques.

### Appendix B: Arduino Codes

In appendix B: The .ino codes for implementation of the pump control mechanism, automated MUX Control, and the overdrive function for the solenoid pinch valves.
